# *Plasmodium falciparum* adapts its investment into replication *versus* transmission according to the host environment

**DOI:** 10.1101/2022.11.29.518379

**Authors:** Abdirahman I. Abdi, Fiona Achcar, Lauriane Sollelis, Joao Luiz Silva-Filho, Kioko Mwikali, Michelle Muthui, Shaban Mwangi, Hannah W. Kimingi, Benedict Orindi, Cheryl Andisi Kivisi, Manon Alkema, Amrita Chandrasekar, Peter C. Bull, Philip Bejon, Katarzyna Modrzynska, Teun Bousema, Matthias Marti

**Affiliations:** KEMRI Wellcome Trust Research Programme, Kilifi, Kenya; Pwani University Biosciences Research Centre, Pwani University, Kilifi, Kenya; Wellcome Center for Integrative Parasitology, Institute of Infection and Immunity, University of Glasgow, Glasgow. Scotland, UK; Institute of Parasitology, Vetsuisse and Medical Faculty, University of Zurich, Zurich, Switzerland; Radboud University Medical Center, Nijmegen, The Netherlands

## Abstract

The malaria parasite life cycle includes asexual replication in human blood, with a proportion of parasites differentiating to gametocytes required for transmission to mosquitoes. Commitment to differentiate into gametocytes, which is marked by activation of the parasite transcription factor *ap2-g*, is known to be influenced by host factors but a comprehensive model remains uncertain. Here we analyze data from 828 children in Kilifi, Kenya with severe, uncomplicated, and asymptomatic malaria infection over 18 years of falling malaria transmission. We examine markers of host immunity and metabolism, and markers of parasite growth and transmission investment. We find that inflammatory responses and reduced plasma lysophosphatidylcholine levels are associated with markers of increased investment in parasite sexual reproduction (i.e., transmission investment) and reduced growth (i.e., asexual replication). This association becomes stronger with falling transmission and suggests that parasites can rapidly respond to the within-host environment, which in turn is subject to changing transmission.

## Introduction

Malaria remains one of the world’s major public health problems. In 2020, an estimated 627’000 deaths and 241 million cases were reported^1^. Around 70% of deaths are in African children under five years of age and are caused by a single parasite species, *Plasmodium falciparum*^1^.

*P. falciparum* has a complex life cycle, involving obligatory transmission through a mosquito vector and asexual replication within erythrocytes of the human host. Between-host transmission requires the formation of gametocytes from asexual blood stage forms, as gametocytes are the only parasite stage to progress the cycle in the mosquito. A series of recent studies has demonstrated that commitment to gametocyte formation (i.e., stage conversion) is epigenetically regulated and occurs via activation of the transcription factor, AP2-G that in turn induces transcription of the first set of gametocyte genes^2,3^.

The parasites that do not convert into gametocytes continue to replicate asexually, contributing to within-host parasite population growth (i.e., parasite burden) and determining *P. falciparum* infection outcome that ranges from asymptomatic infections to severe complications and death^4–6^. Cytoadhesion of infected erythrocytes (IE) to receptors on microvascular endothelium of deep tissues reduces the rate of parasite elimination in the spleen^7,8^, thus supporting the within-host expansion of the parasite population (i.e., parasite burden). As a side effect of this parasite survival strategy, cytoadhesion reduces the diameter of the vascular lumen, thus impairing perfusion and contributing to severe malaria pathology^9–11^. *P. falciparum* erythrocyte membrane protein 1 (PfEMP1), encoded by the *var* multi-gene family, plays a critical role in both pathogenesis (through cytoadhesion)^12,13^ and establishment of chronic infection (through variant switching and immune evasion)^14,15^.

Both *var* gene transcription and stage conversion (and hence *ap2-g* transcription) are subject to within-host environmental pressures such as immunity^16^, febrile temperature^17,18^, and nutritional stress^19^, perhaps via a common epigenetic regulation mechanism^20^. For example, *in vitro* studies revealed that stage conversion can be induced by nutritional depletion such as spent culture media^19,21^ and depletion of Lysophosphatidylcholine (LPC)^22,23^. Recent work from Kenya and Sudan provides some evidence that parasites in low relative to high transmission settings invest more in sexual commitment and less in replication and *vice versa^16^*. Altogether these studies suggest that the parasite can sense and rapidly adapt to its environment *in vitro* and *in vivo*. A family of protein deacetylases called sirtuins are known as signaling proteins linking environmental sensing to various cellular processes via metabolic regulation^24–26^. They do this through epigenetic control of gene expression^26^ and post-translational modification of protein function^25,27^. The *P. falciparum* genome contains two sirtuins (Pfsir2a/b) which have been linked to the control of *var* gene transcription^28,29^, and their expression is influenced by febrile temperature^17^ and low transmission intensity^16^.

Here we investigated the interplay between parasite and host environmental factors governing parasite investment in reproduction (to maximize between-host transmission) *versus* replication (to ensure within-host persistence) *in vivo*. We analyzed samples and clinical data collected from children in Kilifi county, Kenya, over changing malaria transmission intensity between 1994 and 2014. We quantified parasite transcripts for *ap2-g, PfSir2a*, and *var* genes, as well as *Pf*HRP2 protein levels (for parasite biomass) and levels of host inflammatory markers and lipid metabolites. We then integrated these host and parasite-derived parameters to interrogate their dynamics and interactions in the context of changing transmission intensity and immunity.

## Results

### A clinical malaria patient cohort across changing transmission periods in Kilifi, Kenya

The study included samples and clinical data collected from 828 children from Kilifi county, Kenya, over 18 years of changing malaria transmission^30,31^. The study period encompassed three defined transmission phases^31^: pre-decline (1990-2002), decline (2003-2008), and post-decline (2009-2014) (**Fig.1A**). During the study period, a total of 26’564 malaria admissions were recorded at Kilifi county hospital (**Fig.1A, 1B**). While the number of parasite-positive admissions decreased, the mean patient age at admission increased over time^4,6,32^ (**Fig.1A**). For our study, 552 of the admissions were selected to ensure adequate sampling of the transmission periods and clinical phenotypes (**Fig.1C**). 150 patients presented with uncomplicated malaria and 402 with one or a combination of the severe malaria syndromes: impaired consciousness (IC), respiratory distress (RD), and severe malaria anemia (SMA)^33^ (**Fig.1D**). 223 samples from children presenting with mild malaria at outpatient clinics and 53 asymptomatic children from a longitudinal malaria cohort study were added to cover the full range of the possible outcomes of malaria infection (**Fig.1B-E**), bringing the total number included in this study to 828 children. The characteristics of participants and clinical parameters are summarized in **Table S1.** In the subsequent analysis, only clinical cases were considered. Asymptomatic patients were excluded except for analysis about clinical phenotype since asymptomatic sampling was limited to the decline and post-decline periods (**Fig.1C**).

**Figure 1.**
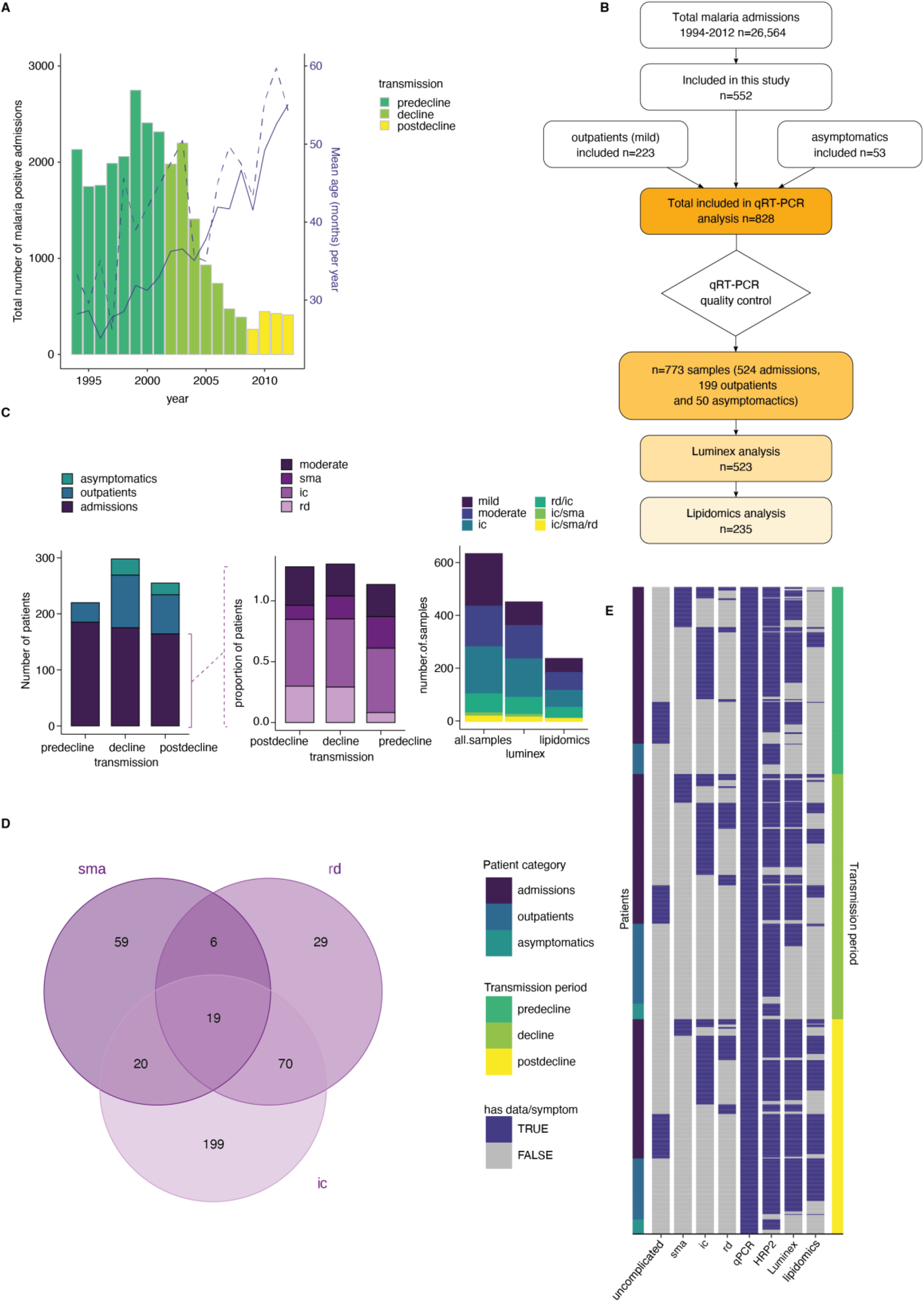
A clincal malaria patient cohort during changing transmission in Kilifi, Kenya. **A**. Total malaria admissions and patient age of the parent cohort. Number of patients per year (grey histogram, left axis). The solid blue line is the average patient age in the parent cohort, the dashed line is the average patient age in this study (both right axis). **B.** Schematic of sample selection for this study. **C.** Clinical presentation of patients selected for this study. Left: all patients, middle: admissions only, right: subset selected for luminex and lipidomics analysis. sma=severe malarial anemia, ic=impaired consiousness, rd=respiratory distress. **D**. Number of patients in this study with different clinical presentations (402 severe cases initially selected). **E.** Overview of the data available for each patient of the study, after excluding samples with *Pfsir2a* and *ap2-g* transcript transcription units greater or equal 32 as described in the methods. Each row is one patient, organised by patient category (left axis) and transmission period (right axis).

### Dynamics of parasite parameters across transmission period and clinical phenotype

First, we analyzed the dynamics of parasite parameters across transmission periods and clinical outcomes. For this purpose, we measured both total parasite biomass based on *Pf*HRP2 levels and peripheral parasitemia based on parasite counts from blood smears. Total parasite biomass decreased with declining transmission (**Fig.2A**). This decrease was significant in the patients presenting with mild malaria at outpatient clinics (**Fig.2A**) which is a more homogenous clinical subgroup as compared to admissions consisting of a range of clinical phenotypes (**Fig.1C-D**).

**Figure 2.**
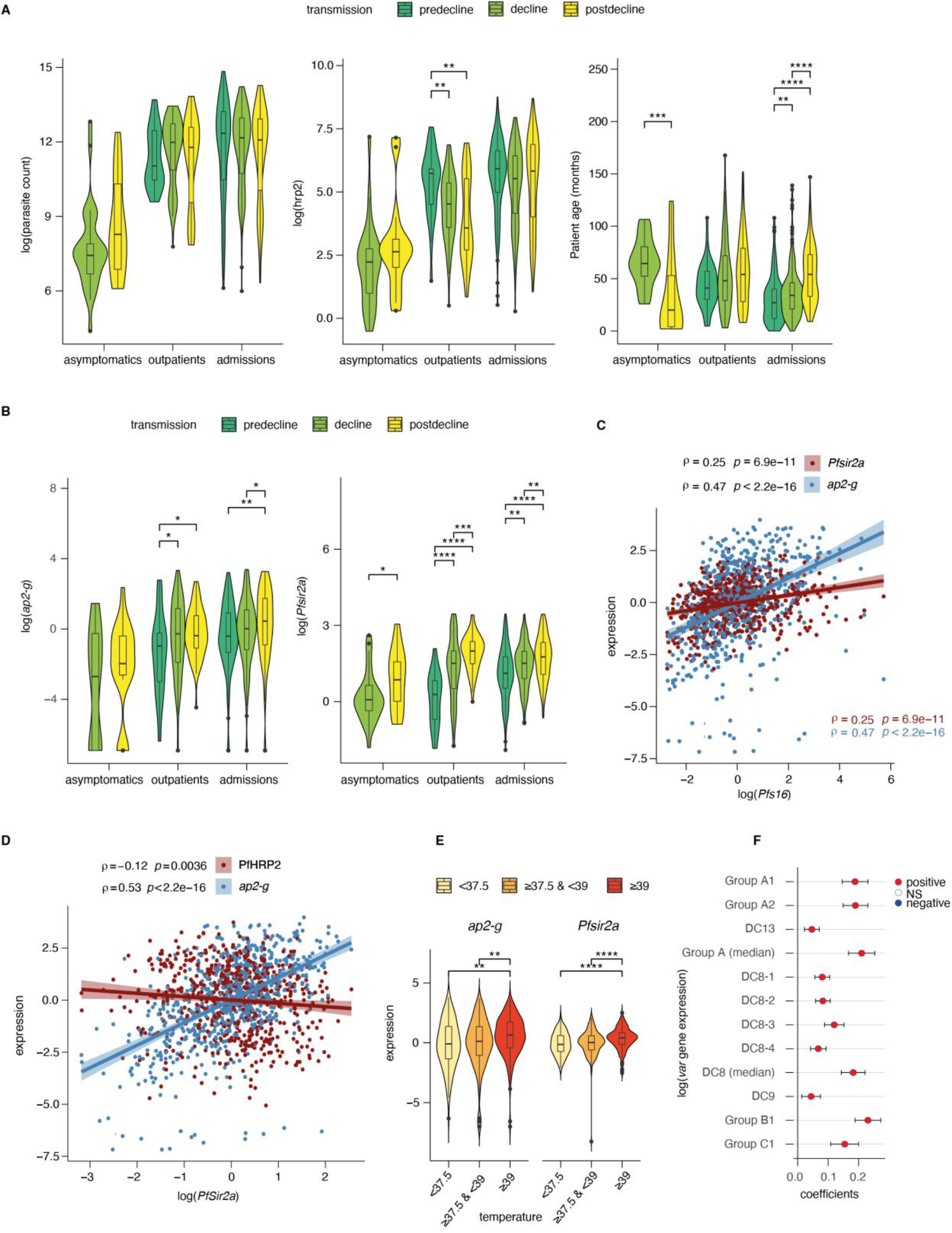
Dynamics of parasite parameters across transmission periods. **A.** Peripheral parasitemia (smear, left), total parasite biomass (*Pf*HRP2, middle) and patient age (right) across patients. **B.***ap2-g* transcript levels (left) and *Pfsir2a* levels (right) across patients. **C.** Spearman’s correlation between *Pfs16* and *ap2-g* (blue) or *PfSir2a* transcription (red) across patients (corrected for transmission). The lines fitted are linear regressions for visualisation only. **D.** Spearman’s correlation between *Pfsir2a* and *ap2-g* transcription (blue) or *Pf*HRP2 levels (red) across patients (corrected for transmission). The lines fitted are linear regressions for visualisation only. **E.***ap2-g* and *Pfsir2a* transcription (corrected for transmission) stratified by patient temperature. **F.** Linear regression of *var* gene transcription levels with *Pfsir2a* levels (adjusted for transmission). 95% confidence intervals are shown. The color indicates whether the relationship is statistically significant (with Benjamini & Hochberg multiple tests correction). Positive correlations in red, negative in blue. In above figures, asymptomatics were excluded in analyses involving transmission period since they are not represented in the pre-decline period. All pairwise statistical tests indicated in the graphs are wilcoxon tests corrected for multiple testing (Benjamini & Hochberg, *=FDR<0.05, **=<0.01, ***=<0.001 and ****=<0.0001).

Parasite samples were subjected to qRT-PCR analysis to quantify transcription of *ap2-g, Pfsir2a*, and *var* gene subgroups relative to two housekeeping genes (fructose biphosphate aldolase and seryl tRNA synthetase)^34,35^. In line with recent findings^16^, *ap2-g* transcription increased significantly with declining malaria transmission (**Fig.2B**). Importantly, *ap2-g* transcription showed a highly significant correlation with transcription levels of the gametocyte marker *Pfs16* (**Fig. 2C**). This association validates *ap2-g* as a proxy for both, stage conversion and gametocyte levels. *Pfsir2a* transcription followed the same trend across transmission periods and was positively associated with *ap2-g* transcription (**Fig.2B,D**). *Pfsir2a* and *ap2-g* transcription also showed a positive association with fever (**Fig. 2E**), suggesting that both factors are sensitive to changes in the host inflammatory response. *Pfsir2a* but not *ap2-g* transcription also showed a significant negative association with *Pf*HRP2 (**Fig.2D**). Given this unexpected observation, we investigated the well-established associations between *Pfsir2a* transcription and *var* gene transcription patterns^29,34^. *Pfsir2a* transcription showed a positive association with global upregulation of *var* gene transcription, particularly with subgroup B (**Fig.2F**). Likewise, transcription of group B *var* subgroup, *Pfsir2a* and *ap2-g* transcription followed a similar pattern in relation to clinical phenotypes (**Fig.S1**). Altogether these data suggest co-regulation of *ap2-g* and *Pfsir2a* and a negative association between *Pfsir2a* and *Pf*HRP2, likely through host factors that are changing with the declining transmission.

### *ap2-g* and *Pfsir2a* transcription is associated with a distinct host inflammation profile

We hypothesized that the observed variation in *ap2-g* and *Pfsir2a* levels across the transmission period and clinical phenotype is due to underlying differences in the host inflammatory response. To test this hypothesis, we quantified 34 inflammatory markers^36^ with Luminex xMAP technology in the plasma of the 523 patients from the outpatient and admissions groups. These patients were selected from the original set of 828 to ensure adequate representation of the transmission periods and clinical phenotypes (including fever), as summarized in **Fig. 1C, E**. For this analysis, all associations were corrected for patient age and *Pf*HRP2 levels as possible confounders.

The markers MCP-1, IL-10, IL-6, IL-1ra were significantly positively correlated with *ap2-g* and *Pfsir2a* transcription (**Fig. 3A and S2**). To cluster the inflammatory markers based on their correlation within the dataset, we used exploratory factor analysis and retained five factors with eigenvalues above 1 (**Fig. S3**). Factor loadings structured the inflammation markers into 5 profiles with distinct inflammatory states (**Fig. 3B**). F1 consists of a mixture of inflammatory markers that support effector Th1/Th2/Th9/Th17 responses (i.e., hyperinflammatory state), F2 represents a Th2 response, F3 represents markers that support follicular helper T cell development and Th17^37^, F4 represents markers of immune paralysis/tissue-injury linked to response to cellular/tissue injury^38^ and F5 represents the inflammasome/Th1 response^39^. F4 showed a significant positive association with *ap2-g* and *Pfsir2a* transcription and fever (**Fig. 3C**). In contrast, F5 showed a negative association with *ap2-g* and fever while F1 was positively associated with fever (**Fig.3C**). In parallel with the observed decrease in *Pf*HRP2 levels (**Fig.2A**), F1 and F5 significantly declined with falling transmission (**Fig.3D**).

**Figure 3.**
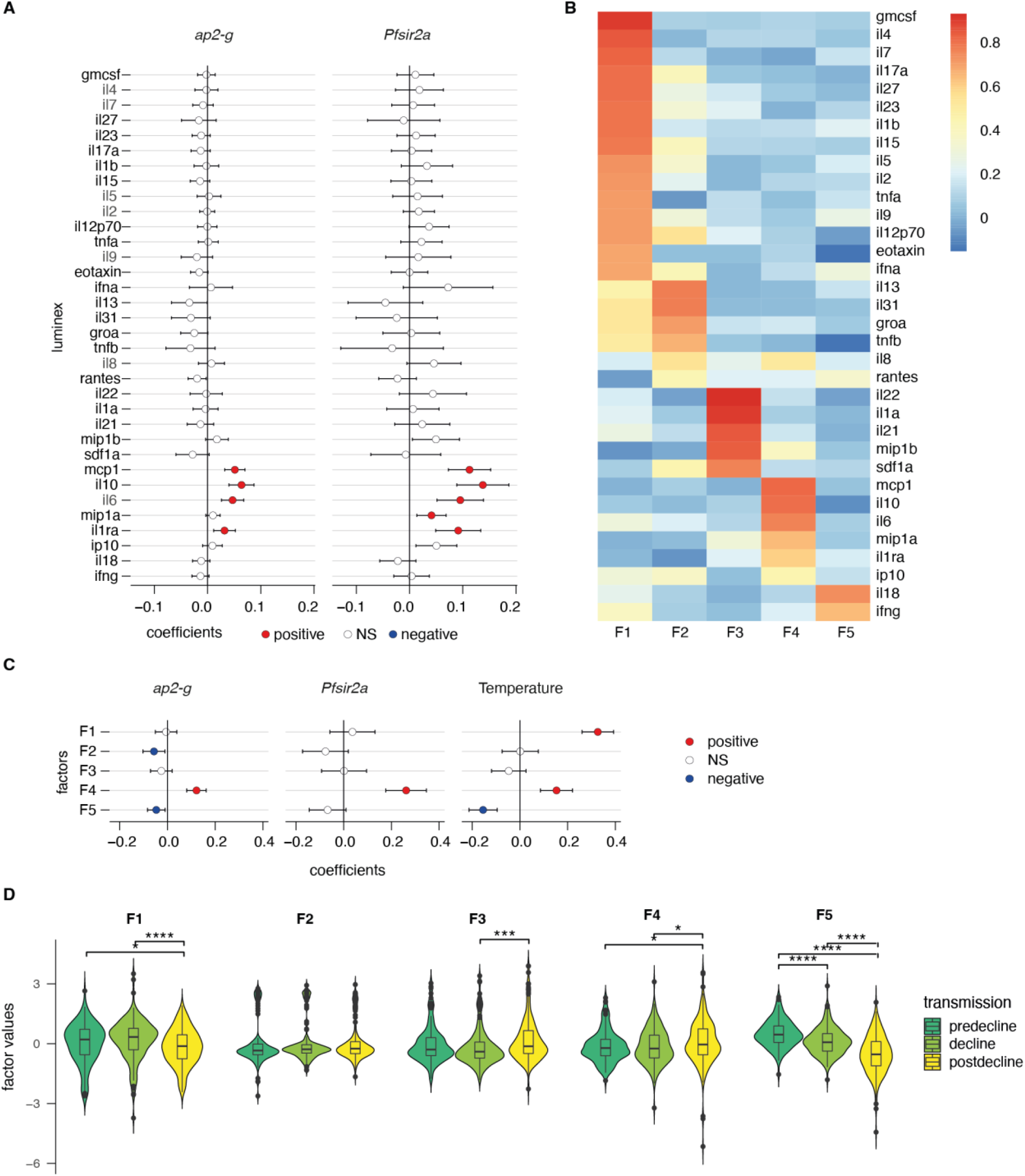
*ap2-g* and *Pfsir2a* transcription levels are associated with the host inflammation profile. **A.** Association of inflammatory markers with *ap2-g* and *Pfsir2a* transcripts, tested using transmission period, age and *Pf*HRP2 adjusted linear regression (*p*-values adjusted for multiple testing using Benjamini & Hochberg multiple tests correction). Plotted is the regression coefficient (estimate) and 95%CI. Above and below zero indicate statistically significant positive (red) and negative association (blue), respectively. **B.** Principal exploratory factor analysis. The figure shows the inflammatory marker loadings on the five factors (F1-F5) identified to have eigenvalue above 1. **C.** Linear regression between inflammatory factors (F1-F5) and *ap2-g* and *Pfsir2a* transcription and patient temperature (adjusted for transmission, *Pf*HRP2 and age). Plotted is the coefficient between the factor and the parameter (estimate) and 95%CI. The association is significant if the correlation FDR < 0.05, in which case the positive associations are marked in red and the negative ones in blue. **D.** Inflammatory factors stratified by transmission period. Pairwise tests are wilcoxon tests (Benjamini & Hochberg, *=FDR<0.05, **=<0.01, ***=<0.001 and ****=<0.0001).

The data support our hypothesis and suggest that the host inflammatory response changes with the falling transmission. Of note, the observed negative association between *Pfsir2a* transcription and *Pf*HRP2 levels appears to be independent of the measured cytokine levels (**Fig.S4**) and is hence likely the result of parasite intrinsic regulation of replication.

### Plasma phospholipids link variation in the host inflammatory profile to *ap2-g* and *Pfsir2a* transcription

We have previously demonstrated *in vitro* that the serum phospholipid LPC serves as a substrate for parasite membrane biosynthesis during asexual replication, and as an environmental factor sensed by the parasite that triggers stage conversion^22^. Plasma LPC is mainly derived from the turnover of phosphatidylcholine (PC) via phospholipase A2, while in the presence of Acyl-CoA the enzyme LPC acyltransferase (LPCAT) can drive the reaction in the other direction^40^. LPC is an inflammatory mediator that boosts type 1 immune response to eliminate pathogens^41,42^. LPC turnover to PC can be triggered by inflammatory responses aimed to repair and restore tissue homeostasis rather than eliminate infection^40^. Here we performed an unbiased lipidomics analysis of plasma from a representative subset of the outpatient and admission patients (**Fig.1B-C,E, S5**) to explore whether the host inflammatory profile modifies the plasma lipid profile and consequently *ap2-g* and *Pfsir2a* transcription levels *in vivo*.

We examined associations between the host inflammatory factors (F1-F5) and the plasma lipidome data. Again, these associations were corrected for transmission period, patient age and *Pf*HRP2 levels. 24 lipid species dominated by phospholipids, showed significant association with the inflammatory factors at a false discovery rate below 0.05 (**Fig.4A**). Similar to the observed associations with *ap2-g* and *Pfsir2a* transcription, cytokines in the F4 and F5 factors showed reciprocal associations with various LPC species and phosphatidylcholine/ethanolamine (PC/PE)(**Fig.4A**): F4 showed negative associations with LPC and positive associations with PC/PE, respectively, and *vice versa* for F5 (**Fig.4A**). The positive association of LPC with the F5 inflammatory factor is consistent with previous findings that identified LPC as an immunomodulator that can enhance IFN-γ production and the activation of the NLRP3 inflammasome, which results in increased levels of cytokines such as IL-1β, IL-18, and IL-33^40–45^ and is necessary for eliminating parasites. Depletion of LPC is also associated with elevated markers of tissue injury (F4), perhaps following uncontrolled parasite growth or maladaptive inflammation. In summary, the association of inflammatory factors with lipids identified LPC, PC and PE species as the most significant ones (**Fig.4A**), in line with their known immunomodulatory role. Importantly, we observed the same pattern in a controlled human infection model where parasite densities were allowed to rise to microscopic levels, both after sporozoite and blood-stage infection (**Fig.4B and S6**)^46,47^. Next, we examined the main lipid species associated with the 5 inflammatory factors with respect to *ap2-g* and *Pfsir2a* transcription. Indeed, LPC species showed a negative association with both *ap2-g* and *Pfsir2a* transcription levels (**Fig.4C-E**). The association was only significant in our data when inflammation is highest (and LPC level lowest), which is at low transmission (i.e., post decline). These data provide *in vivo* evidence for the previously observed link between LPC depletion and *ap2-g* activation and strongly suggest that LPC is both, a key immune modulator and a metabolite whose level is sensed by the parasite. Importantly, the key relationships described in figures 2–4 were independently significant in a structural equation model that examined how host immunity modifies the host-parasite interaction, the within-host environment and parasite investment in transmission or replication (**Table S3**).

**Figure 4.**
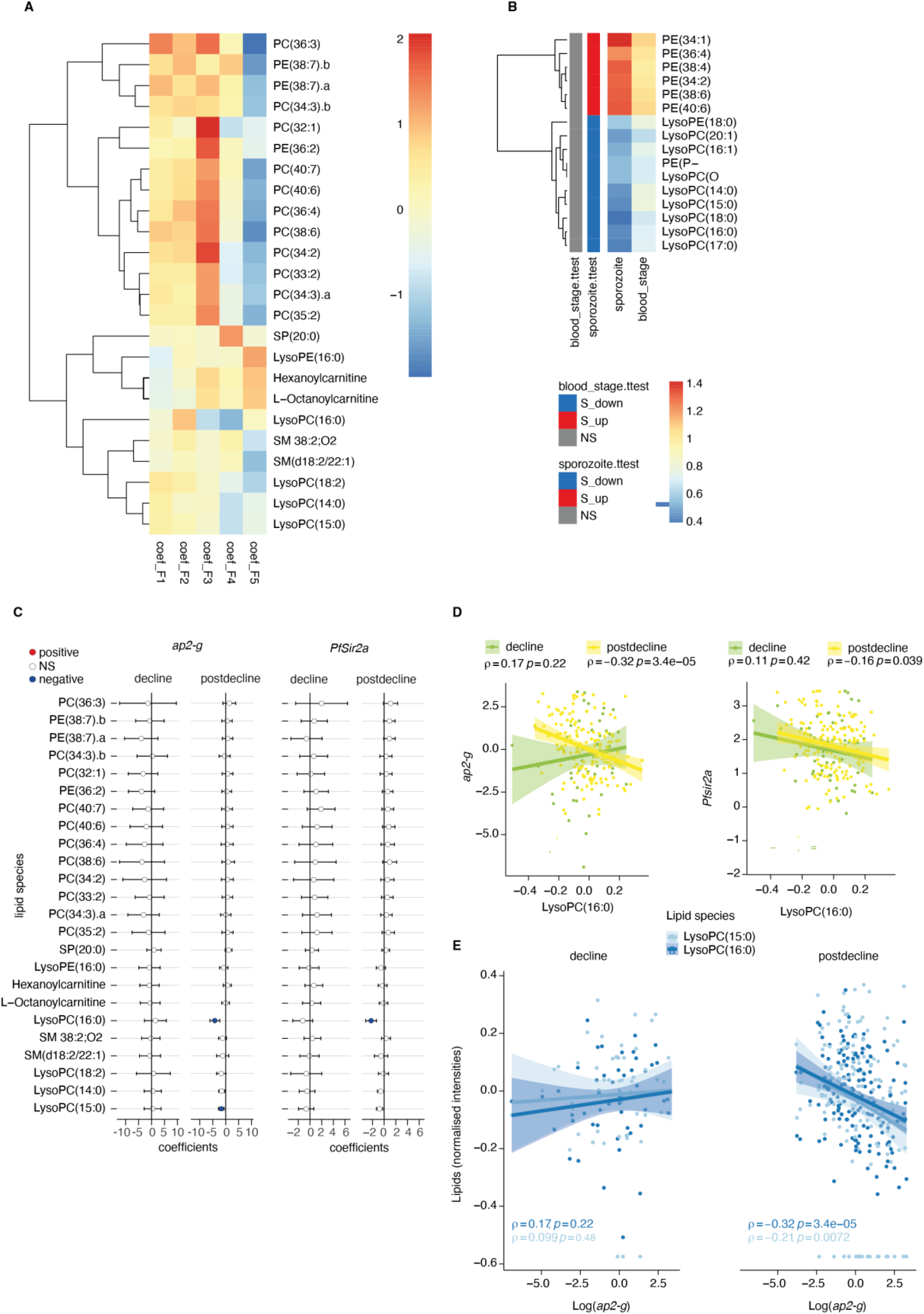
Plasma LPC links host inflammation to *ap2-g* and *Pfsir2a* transcription. **A**. Heatmap of the linear regression coefficients between lipids and inflammatory factors (F1-F5, adjusted for transmission period and corrected for multiple testing). Shown are all lipids that are significantly associated (positive or negative) with factors F1-F5, clustered using R hclust (distance=Euclidean, method=centroid) and that have been manually identified and filtered for peak quality (isotopes and fragments were also filtered out). **B.** Shown are the lipids with significant differences (student’s t-test corrected for multiple testing) between pre- and post-treatment in the controlled human malaria infections (CHMI) for either infection type (blood or sporozoite infection). Plotted is the fold-change post-treatment *vs* pre-treatment. On the left is indicated whether the lipid is significantly increased (red) or decreased (blue) in either type of infection. **C**. Linear regression between the lipids from A and *ap2-g* or *Pfsir2a* transcription levels. Plotted is the coefficient and 95%CI. Blue dots are the statistically significant negative correlations, red are the statically significant positive correlations (FDR<0.05). **D**. Correlation between LPC (16:0) (top) and *ap2-g (top*) or *Pfsir2a* (bottom) transcription (Spearman’s correlations corrected for multiple testing). **E.** Correlations between identified LPCs and *ap2-g* transcription by transmission period (Spearman’s correlations corrected for multiple testing). Note that predecline period is not plotted separately in panels C-E due to insufficient sample numbers for the statistical analysis.

## Discussion

Malaria parasites must adapt to changing environmental conditions across the life cycle in the mammalian and mosquito hosts. Similarly, changing conditions across seasons and transmission settings require both within- and between-host adaptation to optimize survival in the human host *versus* transmission to the next host. First, a recent transcriptomic study from Kenya and Sudan suggested that parasites in low transmission settings (where within-host competition is low) invest more in gametocyte production compared to high transmission settings (where within-host competition is high)^16^. Second, a longitudinal study from Senegal demonstrated that human-to-mosquito transmission efficiency (and gametocyte density) increases when parasite prevalence in the human population decreases, suggesting that parasites can adapt to changes in the environment^48^. However, the within-host mechanisms driving parasite adaptation to the prevailing environment remain unclear.

Here, we analysed parasite and host signatures in the plasma from a large malaria patient cohort over 18 years of declining malaria transmission in Kenya. This investigation allowed us to define some of the within-host environmental factors that change with transmission intensity and consequently influence the parasite decision to invest in reproduction *versus* replication. A major strength of our study is that observations are from a single site and are thus plausibly reflective of transmission-related changes in parasite investments, rather than differences between geographically distinct parasite populations. We show that high transmission is associated with a host immune response that promotes parasite killing without compromising the intrinsic replicative ability of the individual parasite. In contrast, low transmission is associated with a host immune response that increases within-host stressors (fever, nutrient depletion), which trigger higher parasite investment into transmission (see also model in **Fig.5**). Importantly, the observed associations between the parasite parameters *ap2-g*, *Pfsir2a* and host inflammation remain significant if corrected for transmission, but they are strongest at low transmission (i.e., post decline period) when inflammation and the risk of damaging the host are highest.

**Figure 5.**
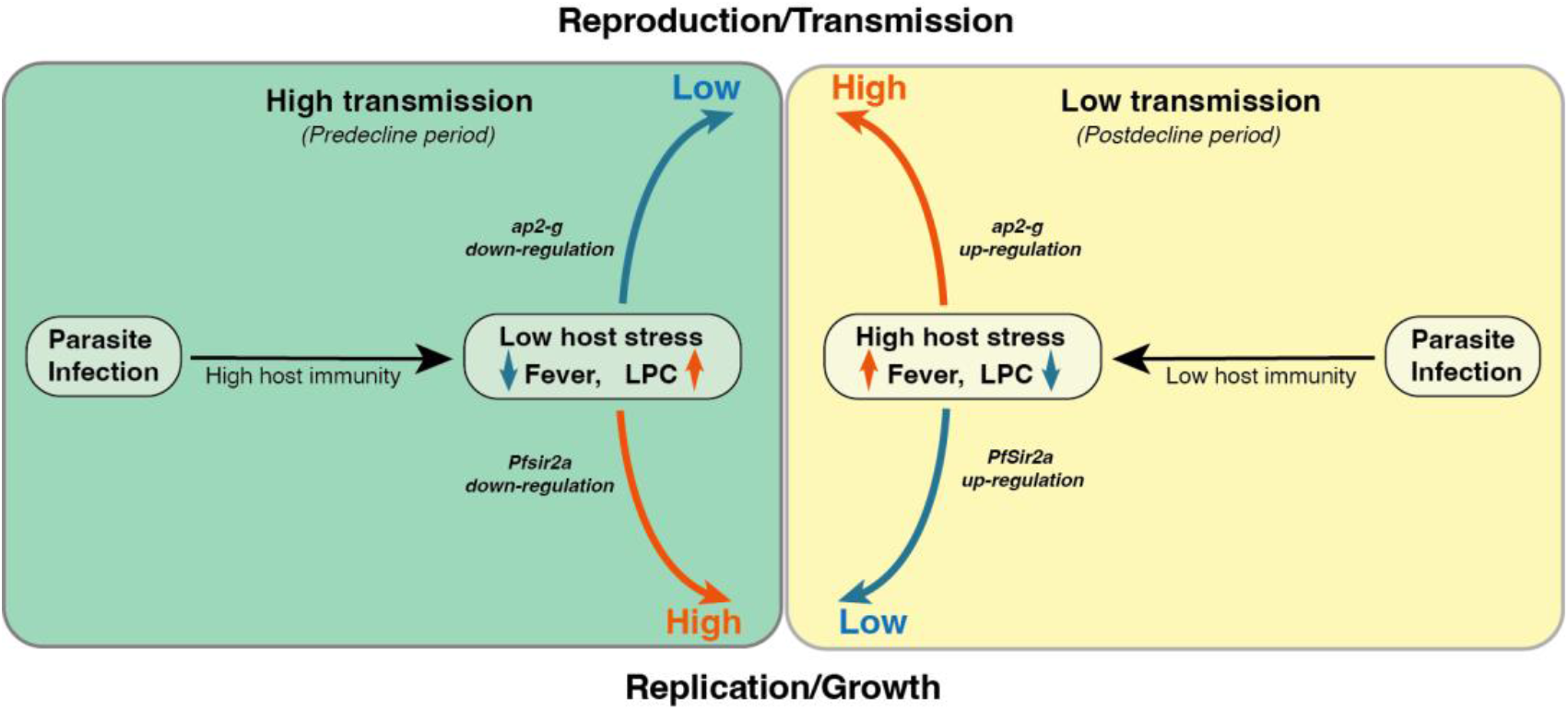
Proposed model on within-host adaptation of the parasite to changing environments. The model is based on the interaction between the different host and parasite parameters described in this study. It proposes that declining transmission reduces host immunity, resulting in increased inflammation (including reduced LPC availability, fever) and susceptibility to clinical symptoms/damage. The altered host response modifies the parasite response during infection, resulting in increased investment in transmission (as indictaed by the elevated *ap2-g* levels) and reduced replication (as indicated by elevated *Pfsir2a* levels and redcued parasite burden/*Pf*HRP2 levels).

At a systemic level, inflammation can influence the within-host environment and modulate parasite investment in replication *versus* reproduction by altering the levels of environmental stressors (e.g., oxidative, thermal, or nutritional stress). Consistent with this hypothesis, we show that a pro-inflammatory response mediated by IFN-γ/IL-18 (F5 in our analysis) promoting pathogen killing^39,49,50^ is negatively associated with *ap2-g* and *Pfsir2a* transcription. In contrast, inflammatory markers that increase within-host environmental stress (e.g., fever) or reflect the extent of host tissue injury and are secreted to heal and restore homeostasis rather than kill pathogens (F4) are positively associated with *ap2-g* and *Pfsir2a* transcription. At a metabolic level, we previously demonstrated that LPC depletion induces *ap2-g* transcription and therefore gametocyte production *in vitro^22^*. A recent study has provided first indications of a possible association between LPC and *ap2-g* levels in a small malaria patient cohort^51^. Here, we reveal that LPC levels are negatively associated with *ap2-g* transcription in patient plasma, thus providing direct evidence for our *in vitro* findings^22^ across a large malaria patient cohort. LPC is an immune effector molecule promoting macrophage polarization to M1 phenotype that induces the secretion of various cytokines such as IFN-γ and IL-1 family (i.e., IL-18) through activation of the NLRP3 inflammasome in endothelial cells and peripheral blood mononuclear cells (PBMCs)^40–45^. Furthermore, LPC is the main component of the oxidized form of LDL (oxLDL) that induces inflammasome-mediated trained immunity in human monocytes^44,45^, resulting in increased responsiveness to LPS re-stimulation. Indeed, we demonstrate that LPC levels are positively associated with IFN-γ/IL-18 levels (Factor 5). These observations are in line with recent data from experimentally infected macaques and malaria patients, where decreased LPC levels were associated with acute *versus* chronic malaria^52^. LPC is also a nutritional resource required by the parasite for replication^22^ and hence scarcity is expected to promote reproduction, as gametocytes require less nutritional resource and therefore provide a better adaptation strategy.

Surprisingly, we also identified a link between *Pfsir2a* transcription, host inflammatory response and parasite biomass (*Pf*HRP2). *Pf*Sir2a belongs to the evolutionarily conserved family of sirtuins that act as environmental sensors to regulate various cellular processes^24,25,53^. In *P. falciparum*, PfSir2a and PfSir2b paralogues cooperate to regulate virulence gene transcription including *var* genes^28,54^. *In vitro* data have also demonstrated that increased PfSir2a levels are associated with reduced parasite replication (i.e., lower merozoite numbers)^55^. We hypothesise that the observed upregulation of *Pfsir2a* transcription in response to inflammation is part of an orchestrated stress response linking replication and antigenic variation (via *Pfsir2a*) to reproduction and transmission (via *ap2-g*), perhaps through a shared epigenetic control mechanism^20^. It is well known that host tolerance to malaria infection reduces with falling transmission^56,57^, as shown by the declining threshold of parasite biomass (*Pf*HRP2) required for clinical malaria. This suggests that parasites have more pronounced harmful consequences on the infected host (i.e., clinical symptoms) in low compared to high transmission settings, perhaps due to increasing host age^58^. Under this scenario, we propose that parasites experience increased within-host stress to which they respond through increased *ap2-g* transcription (to increase reproduction, hence transmission) and increased *Pfsir2a* transcription (to affect antigenic variation and replication, hence the negative association with *Pf*HRP2) – as part of a self-preservation strategy in the face of imminent risk of host death.

In summary, we propose a model where the falling host immunity with declining transmission modifies the predominant host immune response, and consequently, the within-host environment (e.g., LPC availability, fever), resulting in increased investment in transmission (i.e., higher *ap2-g* transcription) and limiting replication (i.e., higher *Pfsir2a* transcription). Our findings provide critical information to accurately model parasite population dynamics. They suggest that parasite populations in elimination scenarios may increase their transmission potential. Understanding how malaria parasites adapt to their environment, for example by increasing investment in transmission stages at low endemicity, is highly relevant for public health. Not only would this affect the timelines for successful elimination, but it would also form an important argument for the deployment of gametocytocidal drugs once transmission has been successfully reduced.

## Materials and Methods

### Study design and participants

Ethical approval was granted by the Scientific Ethics Review Unit of the Kenya Medical Research Institute under the protocol; KEMRI/SERU/3149, and informed consent was obtained from the parents/guardian of the children. The study was conducted at Kilifi county which is a malaria-endemic region along the Kenyan coast. Over the last three decades, Kilifi has experienced changes in the pattern of malaria transmission and clinical presentation spectrum^30–32^. The study included i) children admitted with malaria at Kilifi county hospital (KCH) between 1994-2012 and recruited as part of hospital admission surveillance system, ii) children presenting with mild malaria at out-patient clinic and iii) asymptomatic children which were part of a longitudinal malaria surveillance cohort which were sampled during annual cross-section bleed in 2007 and 2010. Clinical data, parasite isolates and plasma samples collected from the children were used to conduct the study. The selection of sub-samples for quantifying inflammatory markers and lipids was informed by availability of fever data and resource.

### Clinical definitions

Admission to malaria was defined as all hospitalized children with malaria parasitemia. The severe malaria syndromes: severe malarial anemia (SMA), impaired consciousness (IC) and respiratory distress (RD) were defined as haemoglobin <5 g/dl, Blantyre coma score (BCS) <5 and deep breathing, respectively. Malaria admissions that did not present with either of the severe malaria syndromes were defined as moderate malaria. Mild malaria was defined as stable children presenting at outpatient clinic with peripheral parasitemia, and asymptomatic as those with positive malaria (Giemsa smear) but without fever or any other sign(s) of illness. The combination of mild and moderate were referred to as uncomplicated.

### Controlled infection cohort

Malaria naïve volunteers were infected by either bites from 5 *P. falciparum* 3D7–infected mosquitoes (n = 12) or by intravenous injection with approximately 2,800 *P. falciparum* 3D7– infected erythrocytes (n=12); treatment with piperaquine was provided at a parasite density of 5000/mL or on day 8 following blood-stage exposure, respectively ^47^.

### Parasite parameters

Thick and thin blood films were stained with Giemsa and examined for *Plasmodium falciparum* parasites according to standard methods. Data was presented as the number of infected RBCs per 500, 200 or 100 RBC counted. This data was then used to calculate parasitemia per μl of blood using the formula described in “2096-OMS-GMP-SOP-09-20160222_v2.indd (who.int)”. Briefly, parasites/μl= number of parasitized RBCs x number of RBCs per μl /number of RBCs counted or number of parasites counted x number of WBCs per μl/number of WBCs counted. Where data on actual number of RBCs or number of WBCs per μl of blood is not available, 5 million RBC and 8000 WBC per μl of blood was assumed.

### Measurement of cytokine levels in the plasma samples

The selection for this subset was primarily informed by availability of fever data but the transmission period and clinical phenotype were also considered. However, there were more children with fever data record in the post-decline period than pre-decline and decline periods which biased the sampling toward post-decline period. The plasma samples were analyzed using ProcartaPlex Human Cytokine & Chemokine Panel 1A(34plex) [Invitrogen/ThermoFisher Scientific; catalogue # EPX340-12167-901; Lot:188561049] following the manufacturer’s instructions. The following 34 cytokines were measured: GM-CSF, IFN-α, IFN-γ, IL-1α, IL-1β, IL-1RA, IL-2, IL-4, IL-5, IL-6, IL-7, IL-8, IL-9, IL-10, IL-12 (p70), IL-13, IL-15, IL-17A, IL-18, IL-21, IL-22, IL-23, IL-27, IL-31, IP-10 (CXCL10), MCP-1 (CCL2), MIP-1α (CCL3), MIP-1β (CCL4), TNF-α, TNF-β, Eotaxin/CCL11, RANTES, GRO-a, and SDF-1a.

Briefly, 50μl of magnetic beads mix were added into each plate well and the 96-well plate securely placed on a hand-held magnetic plate washer for 2 minutes for the beads to settle. The liquid was then removed by carefully inverting the plate over a waste container while still on the magnet and lightly blotted on absorbent paper towels. The beads were then washed by adding 150μl of 1× wash buffer, left to settle for 2 minutes and the liquid removed as before followed by blotting. This was followed by adding 25μl of Universal Assay Buffer per well and then 25μl of plasma samples and standards into appropriate wells or 25μl of Universal Assay Buffer in blank wells. The plate was covered and shaken on a plate shaker at 500rpm for 30 minutes at room temperature followed by an overnight incubation at 4°C. After the overnight incubation, the plate was shaken on a plate shaker at 500rpm for 30 minutes at room temperature and the beads then washed twice while on a magnetic plate holder as outlined above. The beads were then incubated in the dark with 25μl of detection antibody mixture on a plate shaker at 500rpm for 30 minutes at room temperature followed by two washes as before. A 50μl of Streptavidin-Phycoerythrin (SAPE) solution was then added per well and similarly incubated for 30 minutes on a plate shaker at 500rpm and at room temperature followed by two washes. After the final wash, the beads were resuspended in 120μl of Reading Buffer per well, incubated for 5 minutes on a plate shaker at 500rpm before running on a MAGPIX reader running on MAGPIX xPOTENT 4.2 software (Luminex Corporation). The instrument was set to count 100 beads for each analyte. The analyte concentrations were calculated (via Milliplex Analyst v5.1 [VigeneTech]) from the median fluorescence intensity (MdFI) expressed in pg/mL using the standard curves of each cytokine.

### PfHRP2 ELISA

*Plasmodium falciparum* histidine-rich protein 2 (*Pf*HRP2) was quantified in the malaria acute plasma samples using ELISA as outlined. Nunc MaxiSorp™ flat-bottom 96-well plates (ThermoFisher Scientific) were coated with 100μl/well of the primary/capture antibody [Mouse anti-*Pf*HRP2 monoclonal antibody (MPFM-55A; MyBioscience)] in 1×phosphate buffered saline (PBS) at a titrated final concentration of 0.9μg/ml (stock = 8.53mg/ml; dilution = 1:10,000) and incubated overnight at 4°C. On the following day, the plates were washed thrice with 1×PBS/0.05% Tween-20 (Sigma-Aldrich) using a BioTek ELx405 Select washer (BioTek Instruments, USA) and blotted on absorbent paper to remove residual buffer. These plates were then blocked with 200μl/well of 1×PBS/3% Marvel skimmed milk (Premier Foods; Thame, Oxford) and incubated for 2 hours at room temperature (RT) on a shaker at 500rpm. The plates were then washed thrice as before. After the final wash, plasma samples and standards were then added at 100ul/well and in duplicates. The samples and standards (*Pf*HRP2 Recombinant protein; MBS232321, MyBioscience) had been appropriately diluted in 1×PBS/2% bovine serum albumin (BSA). The samples and standards were incubated for 2 hours at RT on a shaker at 500rpm followed by three washes with 1×PBS/0.05% Tween-20 and blotted dry as before. This was followed by addition of a 100μl/well of the secondary/detection antibody [Mouse anti-*Pf*HRP2 HRP-conjugated antibody (MPFG-55P; MyBioscience) diluted in 1×PBS/2%BSA and at a final titrated concentration of 0.2μg/ml (stock = 1mg/ml; dilution = 1:5,000). The plates were then incubated for 1 hour at RT on a shaker at 500rpm, washed thrice as before and dried on absorbent paper towels. o-Phenylenediamine dihydrochloride (OPD) (ThermoFisher Scientific) substrate was then added at 100μl/well and incubated for 15 minutes for colour development. The reaction was stopped with 50μl/well of 2M sulphuric acid (H_2_SO_4_) and optical densities (OD) read at 490nm with a BioTek Synergy4 reader (BioTek Instruments, USA).

### Parasite transcript quantification using quantative RT-PCR

RNA was obtained from TRIzol™ reagent (Invitrogen, catalog number 15596026) preserved *P. falciparum* positive venous blood samples obtained from the children recruited in the study. RNA was extracted by Chloroform method^59^ and cDNA synthesized using Superscript III kit (Invitrogen, catalog number 18091050) following the manufacturer’s protocol. Parasite gene transcription analysis was carried out through quantitative real-time PCR as described below.

Real-time PCR data was obtained as described^34,60,61^. Four primer pairs targeting DC8 (named dc8-1, dc8-2, dc8-3, dc8-4), one primer pair targeting DC13 (dc13) and two primer pairs targeting the majority of group A *var* genes (gpA1 and gpA2) were used in real-time PCR analysis as described^34^. We also used two primer pairs, b1 and c2, targeting group B and C *var* genes respectively ^62^. Primer pairs targeting *Pfsir2a* and *ap2-g* were also used^34^. Two housekeeping genes, Seryl tRNA synthetase and Fructose bisphosphate aldolase^35,63,64^ were used for relative quantification of the expressed *var* genes, *Pfsir2a* and *ap2-g*. The PCR reaction and cycling conditions were carried out as described^64^ using the Applied Biosystems 7500 Real-time PCR system. We set the cycle threshold (Ct) at 0.025. Controls with no template were included at the end of each batch of 22 samples per primer pairs and the melt-curves analysed for non-specific amplification. The *var* gene “transcript quantity” was determined relative to the mean transcript of the two housekeeping genes, Sery tRNA synthetase and Fructose biphosphate aldolase as decribed^64^. For each test primer, the ΔCt was calculated relative to the average Ct of the two housekeeping genes which was then transformed to arbitrary transcript unit (Tu_s_) using the formula (Tus = 2^(5-Δct)^) as described^64^. We assigned a zero Tus value if a reaction did not result in detectable amplification after 40 cycles of amplification, i.e., if the Ct value was undetermined.

### Lipidomics analysis

Serum samples were preserved at −80°C until extraction with the chloroform/methanol method. 25μL of serum were extracted with 1 mL of the extraction solvent chloroform/methanol/water (1:3:1 ratio), the tubes rocked for 10 min at 4°C and centrifuged for 3 min at 13’000g. Supernatant were collected and stored at −80°C in glass tubes until analysis.

Sample vials were placed in the autosampler tray in random order and kept at 5°C. Separation was performed using a Dionex UltiMate 3000 RSLC system (Thermo Scientific, Hemel Hempstead) by injection of 10 μl sample onto a silica gel column (150 mm × 3 mm × 3 μm; HiChrom, Reading, UK) used in hydrophilic interaction chromatography (HILIC) mode held at 30°C ^65^. Two solvents were used: solvent A [20% isopropyl alcohol (IPA) in acetonitrile] and solvent B [20% IPA in ammonium formate (20 mM)]. Elution was achieved using the following gradient at 0.3 ml/min: 0–1 min 8% B, 5 min 9% B, 10 min 20% B, 16 min 25% B, 23 min 35% B, and 26–40 min 8% B. Detection of lipids was performed in a Thermo Orbitrap Fusion mass spectrometer (Thermo Fisher Scientific Inc., Hemel Hempstead, UK) in polarity switching mode. The instrument was calibrated according to the manufacturer’s specifications to give an rms mass error <1 ppm. The following electrospray ionization settings were used: source voltage, ±4.30 kV; capillary temp, 325°C; sheath gas flow, 40 arbitrary units (AU); auxiliary gas flow, 5 AU; sweep gas flow, 1 AU. All LC-MS spectra were recorded in the range 100–1,200 at 120,000 resolutions (FWHM at m/z 500).

### Data preprocessing

The raw data was converted to mzML files using proteowizard (v 3.0.9706 (2016-5-12)). These files were then analysed using R (v 4.2.1) libraries xcms (v 3.14.1) and mzmatch 2 (v 1.0 - 4) for peak picking, alignment, filtering and annotations^66–68^. Batch correction was applied as in (https://www.mdpi.com/2218-1989/10/6/241/htm), the data was then checked using PCA calculated using the R function prcomp (see supplementary figure 4B). Data was then range normalised and logged transformed using MetaboanalystR (v3.1.0). The CHMI lipidomics data was analysed the same way but did not require batch correction as the samples were run in one batch.

### Statistical analysis

All data were analyzed using R (v4.2.1). We normalized non-normally distributed variables by log transformation.

#### qRT-PCR

Zeros in qRT-PCR values were replaced by 0.001 (value before log transformation as the smallest measured value is about 0.0017). The median transcript units from qRT-PCR were calculated as follows: DC8 median from four primer pairs used (DC8-1, DC8-2, DC8-3 and DC8-4) and group A median from three primer pairs (gpA1, gpA2 and dc13). Samples for which *ap2-g* or *pfsir2a* arbitrary transcript unit was greater or equal to 32 (that is the transcript quantity of the reference genes based on the formula (Tus = 2^(5-Δct)^)^64^) were deemed unreliable and excluded from the analysis that went into generating figures 2–4. Comparison between two groups was done using two-sided wilcoxon test. All correlations were conducted using Spearman’s rank correlation coefficient test. All forest plots were done using linear regressions adjusted for transmission period, *Pf*HRP2 and age of the patient (see figure legends) using R function lm. All multiple test corrections were done using Benjamini & Hochberg multiple test (using R function p.adjust).

#### Principal factor analysis

A measurement model (i.e., factor analytic model) was fitted to summarise the 34 analytes into fewer variables called factors. An exploratory factor analysis (EFA) was performed to explore the factor structure underlying the 34 analytes. Factors were retained based on the Kaiser’s ‘eigenvalue rule’ of retaining eigenvalues larger than 1. In addition, we also considered the scree plot, parallel analysis, fit statistics and interpretability of the model/factors. This analysis resulted in the cytokine data being reduced to 5 factors. This analysis was done using the R “psych” library (v 2.1.9) available at https://CRAN.R-project.org/package=psych. The 34 analytes were individually linearly regressed to *ap2-g* or *Pfsir2* transcript levels with transmission, *Pf*HRP2 and age correction (model: analyte ~ transmission+*pf*HRP2+age). Then each factor was analyzed the same way.

#### Lipidomics analysis

The preprocessed lipidomics data was tested using transmission period, *Pf*HRP2 and age adjusted linear regression with any of the 5 factors. All m/z with a significant false discovery rate with any of the factors were then manually checked for peak quality and identified masses on mass and retention time^69^. The remaining identified lipids were then checked for relationship with *ap2-g* and *Pfsir2a* transcription levels (linear regression adjusted for transmission period, HRP and age, see methods above). The CHMI lipidomics data was analyzed the same way but the peaks retained were those significantly different pre and post treatment in either type of infection (student’s t-tests corrected for multiple testing).

#### Figures

All heatmaps were done using the R library pheatmap (v 1.0.12) available at https://CRAN.R-project.org/package=pheatmap, and all other plots using the R libraries ggplot2 (v 3.3.5) and ggpubr (v 0.4.0) available at https://CRAN.R-project.org/package=ggpubr.

## Data availability

Raw data and script for all the analyses in this manuscript are available at https://doi.org/10.7910/DVN/BXXVRY. Raw mass spectrometry files for all lipidomics data sets are currently being submitted to MetaboLights.

## Acknowledgements

This work was supported by Wolfson Merit Royal Society Award (to M.M.), Wellcome Trust Investigator Award 110166 (to F.A., L.S., J.L.S.F. and M.M.) and Wellcome Trust Center Award 104111 (to F F.A., L.S., J.L.S.F. and M.M.), Glasgow-Radboud collaborative grant (to M.A., T.B. and M.M.), European Research Council (ERC) Consolidator Grant (to T.B., ERC-CoG 864180 QUANTUM), Wellcome Award 209289/Z/17/Z (to A.A.) and a core Wellcome award to KEMRI-Wellcome Trust (203077/Z/16/Z). This paper was published with permission of the director of Kenya Medical Research Institute.

## Supplementary figures and tables

**Figure S1.**
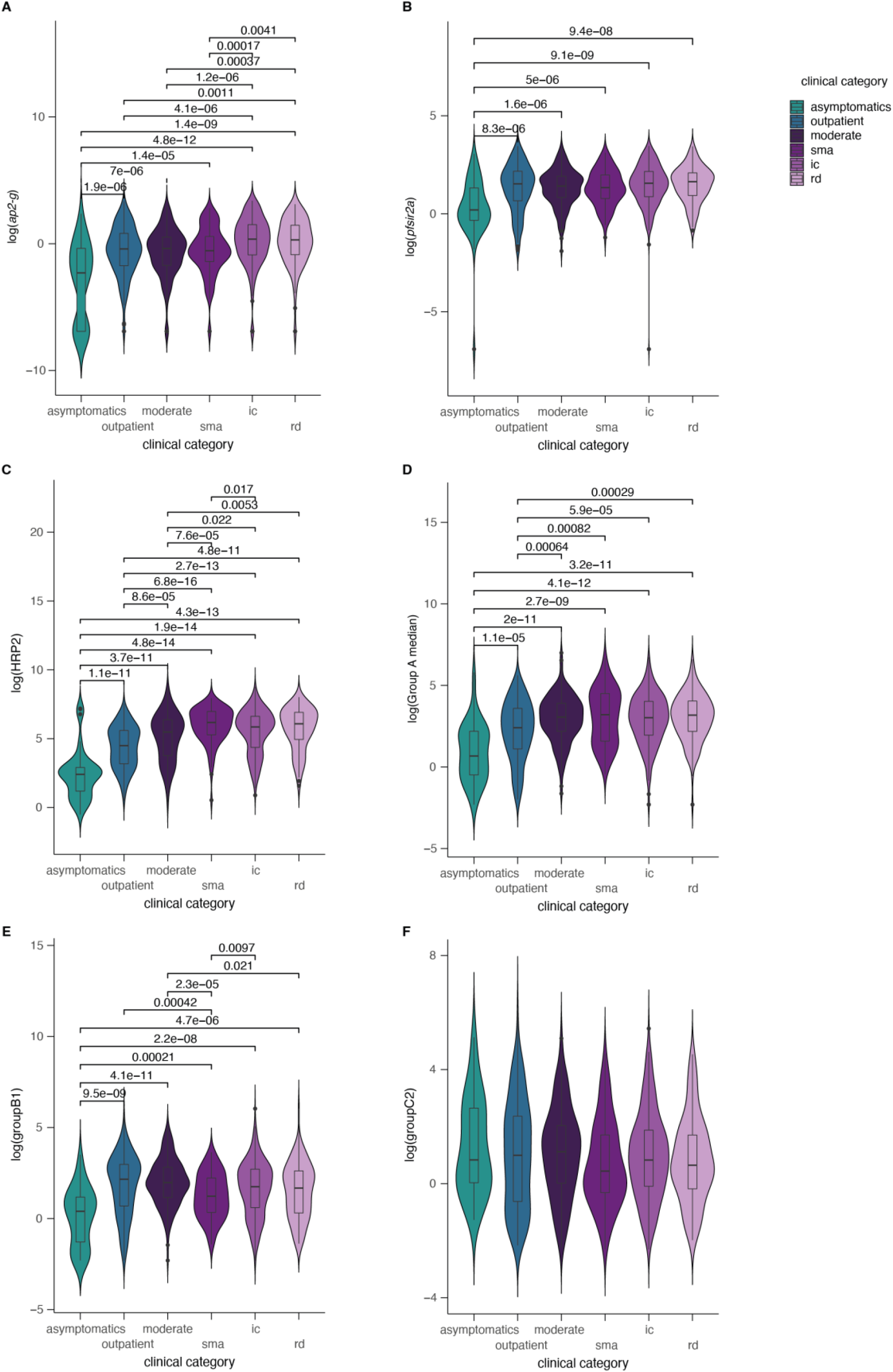
Parasite parameters stratified by clinical categories. *ap2-g, Pfsir2a, var* gene transcription and *Pf*HRP2 levels stratified by clinical categories. Significant wilcoxon test *p*-values (corrected for multiple testing) marked with *<0.05, **<0.01, ***<0.001, ****<0.0001.

**Figure S2.**
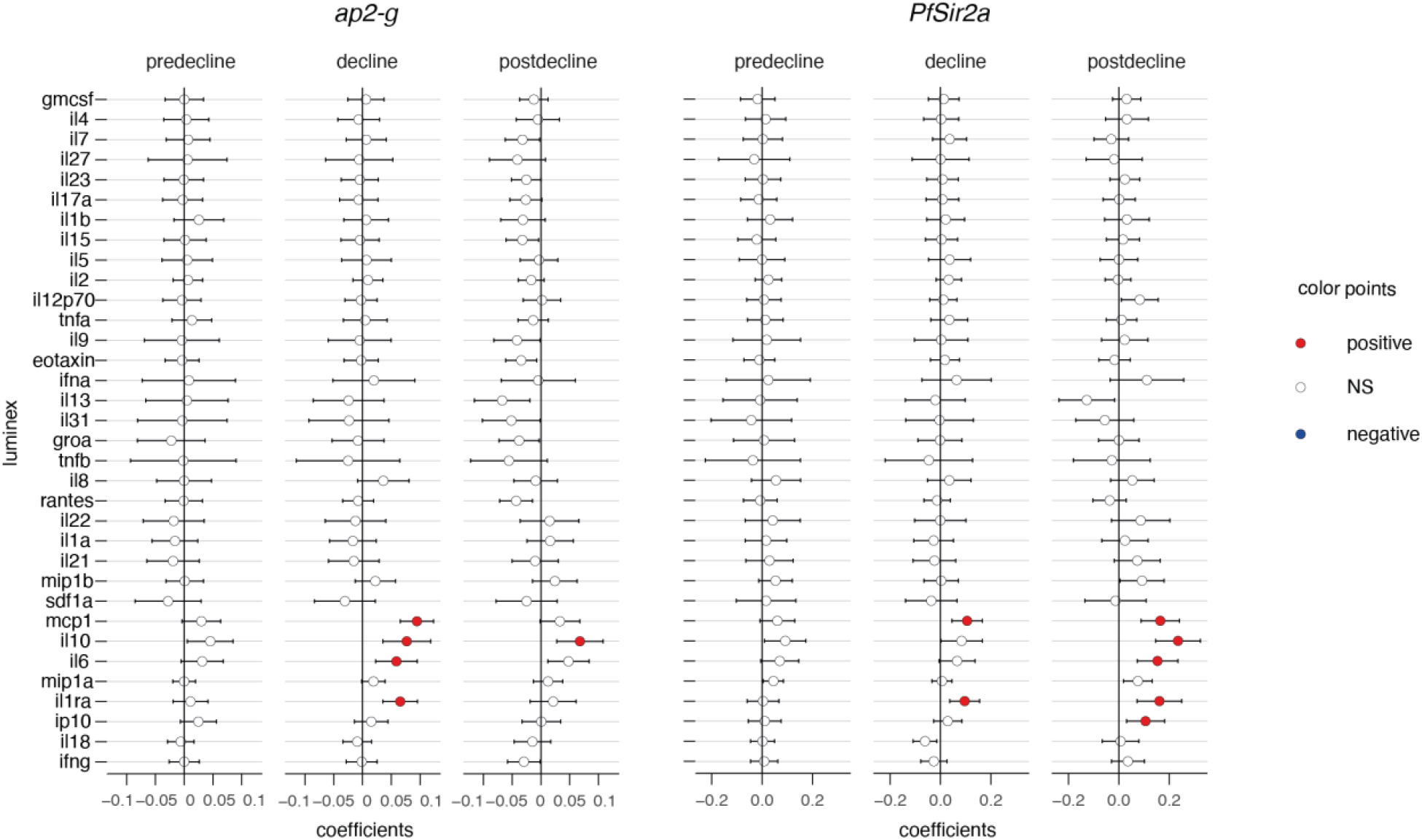
Inflammatory markers stratified by transmission period. Linear regressions between *ap2-g* (**A**) *or Pfsir2a* (**B**) transcription and luminex markers as per figure 3A, stratified by transmission period.

**Figure S3.**
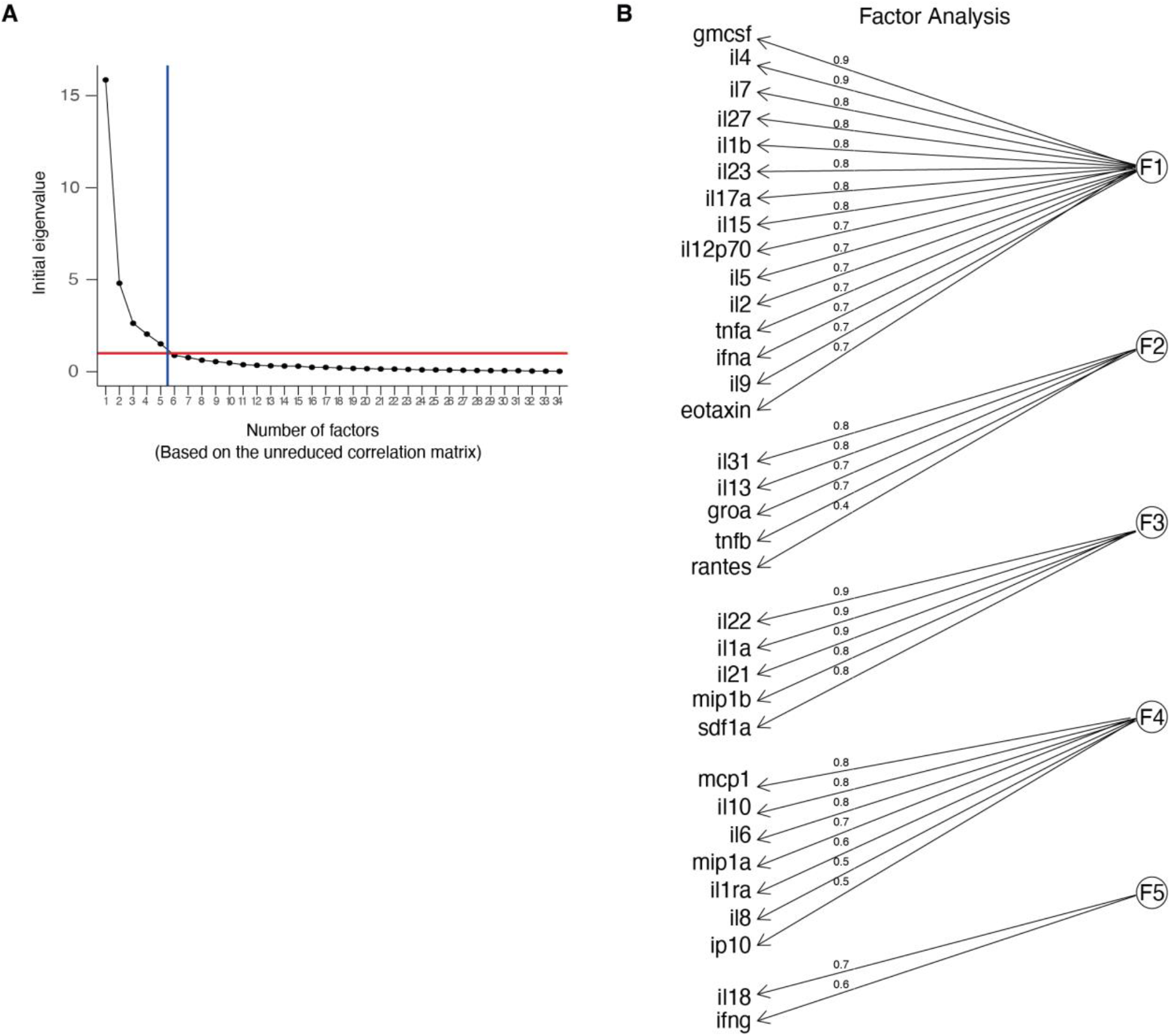
Factor loadings. **A.** Scree plot showing the eigen values *vs* the number of factors (factor analysis of the luminex data). **B.** Major loadings of each factor calculated by factor analysis of the luminex data (values>=0.3). Loading values are noted on the edges.

**Figure S4.**
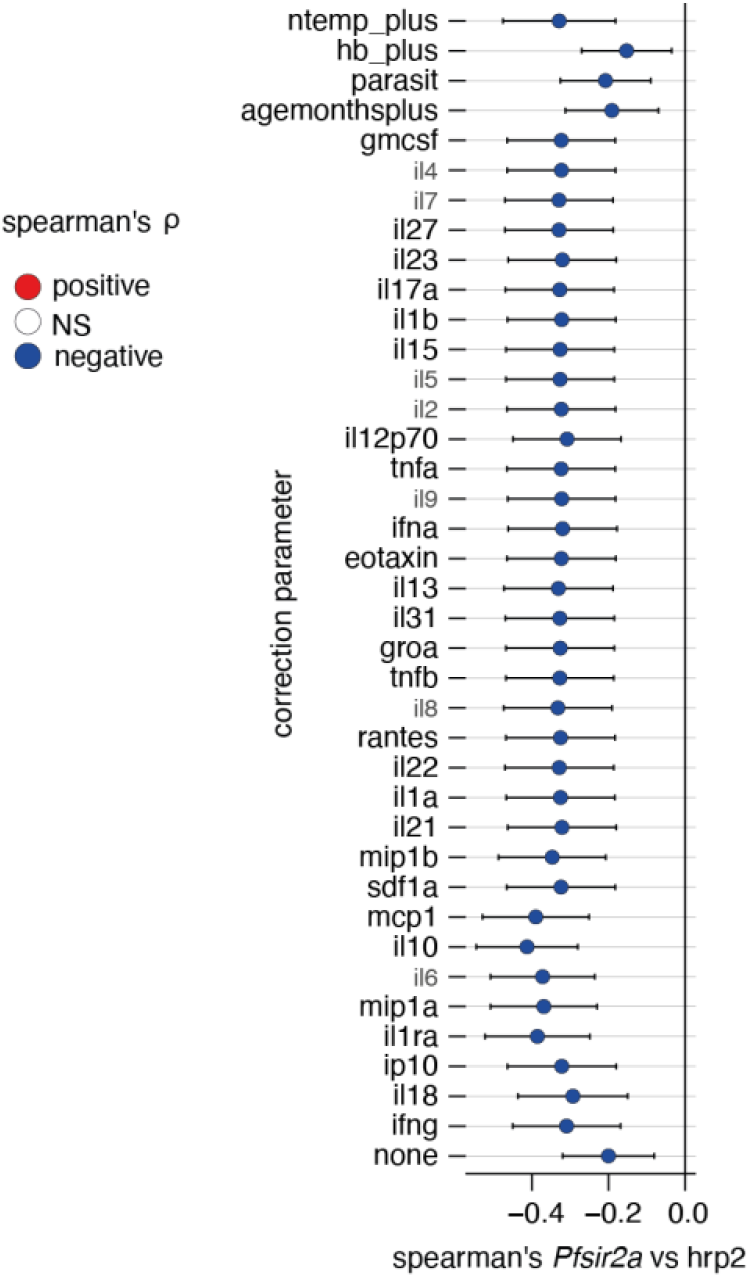
Correcting *Pf*HRP2 vs *Pfsir2a* associations for external factors. Plotted is the linear regression coefficient (estimate) and 95%CI. Blue indicates a significant negative correlation between *Pf*HRP2 levels and *Pfsir2a* transcription with the additional correction indicated on the left.

**Figure S5.**
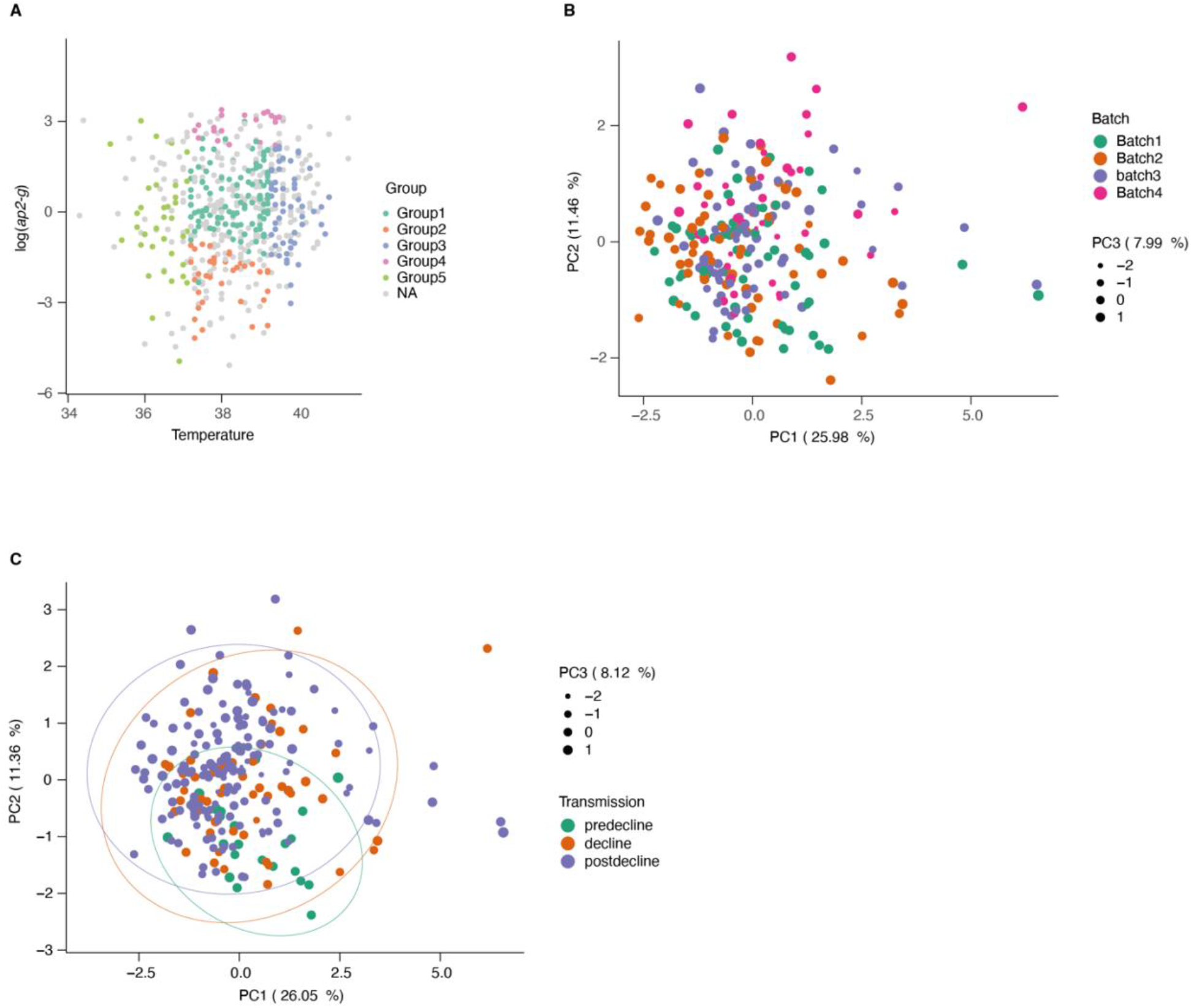
Sample subsetting and batch correction for lipidomics data. **A.** Patient temperature, *ap2-g* transcription level and disease type were used to subset samples for metabolomics. This resulted in 5 groups from severe disease categories and matching mild cases (outpatients, in grey). **B.** PCA of the lipidomics data colored by batch number (post batch correction)**. C.** PCA of the lipidomics data colored by transmission period.

**Figure S6.**
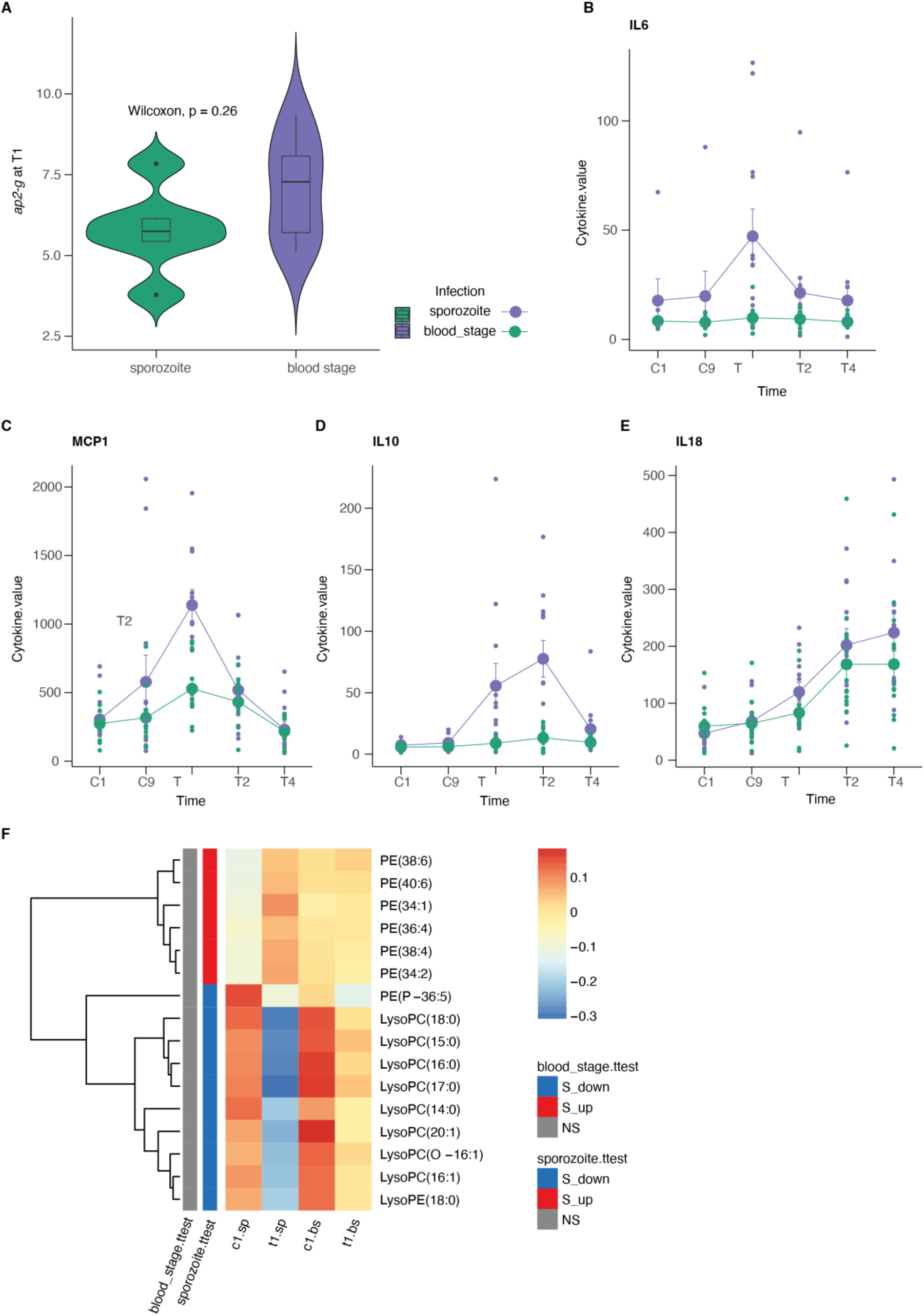
CHMI data. **A.** *ap2-g* transcription measured on day 1 of treatment (T1) and stratified by type of infection. **B-E**: Average (and standard deviation) cytokine levels during the experiment per infection type. Shown are those markers shared with the luminex analysis of the Kilifi cohort. **F.** Average normalized lipid levels are significantly different pre and post treatment (sp=sporozoite infection, bs=blood stage infection). (C=days post infection, T=days post treatment). On the left is indicated whether the lipid is significantly increased (red) or decreased (blue) in either type of infection.

**Table S1.** Associations between parasite parameters *ap2-g, Pfsir2-a* and *Pf*HRP2 and clinical parameters.

**Table S2.** Associations between parasite parameters *ap2-g, Pfsir2-a* and *Pf*HRP2, host luminex markers and lipidomics data.

**Table S3.** Structural equation model (SEM). The model assumes that pre-existing host immunity affects the interaction between host (i.e., altered within host environment including inflammatory response. fever, nutritional resource availability) and parasite (i.e., altered investment in reproduction *vs* replication). Significant *p*-values are highlighted in bold and negative and positive estimates of associations are highlighted in blue and red, respectively.

